# Isoform-Specific Gene Regulation by Progesterone Receptors Drives Divergent Phenotypes in Breast Cancer Cells

**DOI:** 10.1101/2025.05.19.654935

**Authors:** Noelle E. Gillis, Thu H. Truong, Caroline H. Diep, Angela Spartz, Julie H. Ostrander, Carol A. Lange

## Abstract

Exposure to progesterone is a recognized risk factor for breast cancer, and *PGR* polymorphisms are associated with various malignancies. Two progesterone receptor (PR) isoforms, full length PR-B and truncated PR-A, are expressed from the *PGR* gene in breast tissue and play crucial roles in normal physiology and breast cancer progression. An imbalance in the expression ratio of these isoforms, favoring increased levels of PR-A, is common in breast cancer and is associated with resistance to tamoxifen in luminal A-type tumors. Notably, PRs have recently been implicated in promoting endocrine resistance and driving the expansion of cancer stem-like cell (CSC) populations. Despite this insight, the isoform-specific molecular and epigenetic mechanisms underlying PR action in estrogen receptor positive (ER+) breast cancers remain understudied. Phenotypic studies of T47D cell lines that express exclusively PR-A or PR-B showed that PR isoforms regulate divergent cell fates. PR-B-expressing cells have a higher proliferation rate, while PR-A-expressing cells produce more mammospheres. We profiled progesterone-driven gene expression in cells grown in both adherent (2D) and mammosphere (3D) growth conditions and found differential gene regulation by PR-A and PR-B that is consistent with the observed divergent phenotypes. Only the PR-A-driven gene signature of ER+ breast cancer cells maintained as non-adherent mammospheres robustly predicted poor clinical outcome in the METABRIC data set. We then performed CUT&RUN to identify the genomic binding patterns unique to each PR isoform and their suite of target genes. Our findings indicate that PR-A acts as a regulator of the cell cycle, while PR-B plays a pivotal role in metabolism and intracellular signaling. Our genomic profiling of PRs in this model system has unveiled novel isoform-specific functions of PR. This work has shifted our prior understanding of the role of PRs in gene regulation, offering potential insights for therapeutic interventions in ER+ breast cancer.

## INTRODUCTION

Estrogen receptor-positive (ER+) luminal breast cancers account for approximately 80% of all breast cancer cases and are primarily driven by estrogen (E2) signaling. While endocrine therapies targeting ER have significantly improved patient outcomes, up to 40% of ER+ breast tumors develop *de novo* or acquired resistance, ultimately leading to disease progression and poor prognosis. Notably, the majority of relapsed tumors retain steroid hormone receptor (SR) expression (1), suggesting that alternative SR-mediated pathways contribute to therapy resistance. While extensive research has explored mechanisms of ER “escape,” such as ligand-independent activation and *ESR1* mutations (2,3), the role of progesterone receptor (PR) in therapy resistance and metastasis remains underappreciated.

PR is traditionally viewed as a downstream effector of ER signaling, and total PR protein expression serves as a clinical marker of ER activity and responsiveness to endocrine therapy (4). However, increasing evidence suggests that PR is an active transcriptional regulator that co-governs gene expression with ER, fundamentally reshaping the breast cancer transcriptome (5-7). PR is expressed as two isoforms—PR-A and PR-B—that have distinct transcriptional activities (8), yet their individual contributions to tumor biology remain poorly defined. Our previous work demonstrated that phosphorylated PRs (p-PRs) promote the expansion of breast cancer stem-like cell (CSC) populations and drive endocrine resistance (9-11). These findings underscore the need to dissect isoform-specific PR functions at the genomic level and their impact on tumor progression and therapy response.

Epidemiological studies have linked progesterone (P4) exposure to an increased risk of breast cancer (12-17), and genetic polymorphisms in the *PGR* gene have been associated with higher susceptibility to hormone-driven cancers (18-22). In normal breast epithelia, PR-A and PR-B are expressed in a balanced ratio and function cooperatively as A:B heterodimers (23). However, in breast cancer, this equilibrium is frequently disrupted leading to isoform dominance and a prevalence of homodimers, resulting in altered transcriptional outcomes (24,25). Notably, an elevated PR-A:PR-B ratio is often observed in luminal A-type ER+ tumors and has been associated with tamoxifen resistance and poor clinical outcomes (26). Within a breast tumor, this elevated PR-A:PR-B ratio occurs across a heterogeneous population of cells, some of which express exclusively PR-A (27). Despite these observations, most molecular studies consider PR as a singular entity, overlooking the distinct and potentially opposing functions of PR-A and PR-B.

A major challenge in the field has been the inability to reliably distinguish the isoform-specific roles of PR-A and PR-B in cell line models and in patient samples. This gap in knowledge is particularly relevant given that PRs undergo extensive post-translational modifications, including phosphorylation, which modulate their activity in response to oncogenic signaling pathways (28). Understanding how PR-A and PR-B differentially contribute to endocrine resistance and tumor heterogeneity is critical for refining therapeutic strategies targeting PR in ER+ breast cancer.

To address this unmet need, we leverage an isoform-specific model system in which T47D cells express either PR-A or PR-B exclusively (10). By integrating genome-wide PR binding data (CUT&RUN) with transcriptomic analyses (RNA-seq) and phenotypic studies we systematically define the distinct gene regulatory networks governed by each PR isoform. Our study reveals novel insights into the divergent roles of PR-A and PR-B in shaping the breast cancer transcriptome, influencing cell fate, and regulating phenotypic characteristics. By uncovering these isoform-specific functions, we provide a foundation for the development of more precise approaches to targeting PR signaling in ER+ breast cancer.

## RESULTS

### PR-A and PR-B exhibit distinct phenotypic characteristics in vitro

For these studies, we employed PR-null and PR+ variants of ER+ T47D breast cancer models. Each PR isoform was stably re-expressed in naturally occurring PR-null (T47D-Y) cells, as first described by Sartorius et al (29). To demonstrate exclusive expression of PR-A and PR-B isoforms in our modified T47D cell line models, we assessed total and phosphorylated (Ser294) PR-A and PR-B protein levels by Western blot (**Figure 1A**). In the absence of estrogen, T47Dco (29) cells express roughly equal levels of endogenous total PR-A and PR-B in 2D conditions that are phosphorylated upon treatment with R5020. In 3D conditions, PR expression decreases with R5020 treatment, likely due to increased turnover of the liganded receptor (30). PR-A is robustly expressed in T47D PR-A cells grown in 2D and 3D conditions and is phosphorylated in response to R5020 treatment. PR-B is expressed in both 2D and 3D conditions in T47D PR-B cells, with a slight decrease in total PR-B levels in 3D after R5020 treatment.

**Figure 1.**
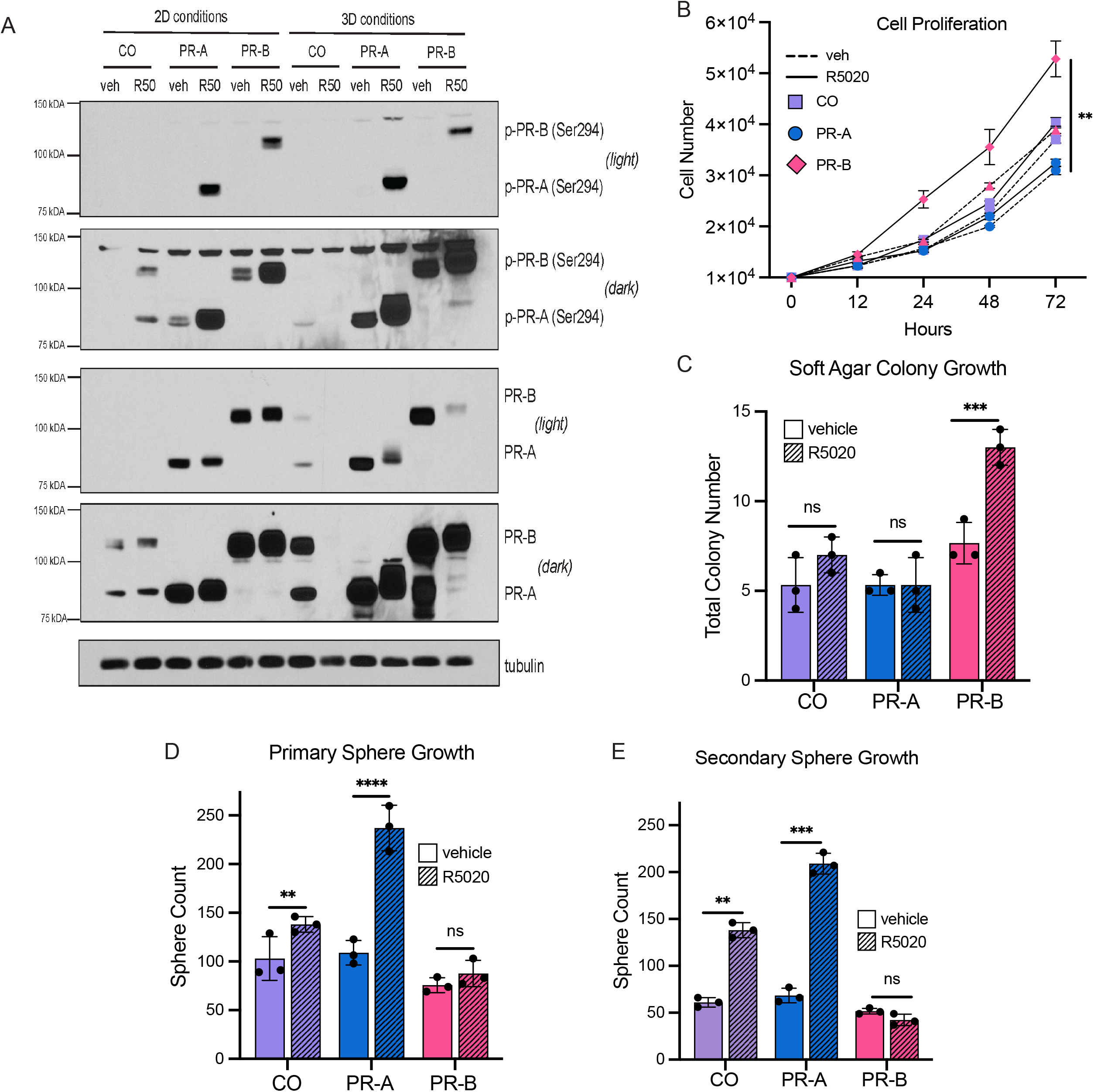
PR-A and PR-B expressing cells exhibit divergent phenotypic characteristics in vitro. A. Western blot shows protein expression of total PR and phosphorylated PR (p-PR) in T47D CO, PR-A, and PR-B expressing cell lines in 2D and 3D growth conditions. B. Cell growth over 72 hours was measured in T47D CO, PR-A, and PR-B expressing cell lines with and without 10 nM R5020 treatment (n =3). C. Quantification of colony formation in soft agar in T47D CO, PR-A, and PR-B expressing cell lines with and without 10 nM R5020 treatment (n = 3). D. Primary sphere counts of T47D CO, PR-A, and PR-B expressing cell lines with and without 10 nM R5020 in the media (n = 3). E. Secondary sphere counts of T47D CO, PR-A, and PR-B expressing cell lines with and without 10 nM R5020 in the media (n = 3).

Cell proliferation assays, reflective of growth in 2D conditions, revealed that PR-A and PR-B differentially regulate cell growth in response to R5020 treatment, with PR-B-expressing cells exhibiting increased proliferation over 72 hours (**Figure 1B**) compared to T47Dco and PR-A-expressing cells. In soft agar colony formation assays, PR-B-expressing cells formed significantly more colonies than T47Dco and PR-A-expressing cells, suggesting an increased anchorage-independent growth potential (**Figure 1C**). Primary and secondary sphere formation assays, where cells are grown in 3D conditions, demonstrated that PR-A-expressing cells had a greater capacity for mammosphere formation with R5020 treatment (**Figure 1D–E**). This indicates a potentially dominant role for PR-A in regulating breast cancer cell stem-like properties.

### PR-A and PR-B regulate distinct transcriptional programs

To determine how PR isoforms influence differential gene expression in response to progesterone, we performed RNA-seq on PR-A and PR-B expressing T47D cells treated with R5020 in both adherent (2D) and mammosphere (3D) conditions. Principal component analysis (PCA) revealed that both growth condition and R5020 treatment were major drivers of transcriptional differences between samples (**Figure 2A**). Unsupervised clustering of all differentially expressed genes further demonstrated distinct transcriptional responses between PR-A and PR-B expressing cells, with notable differences between 2D and 3D culture conditions (**Figure 2B**).

**Figure 2.**
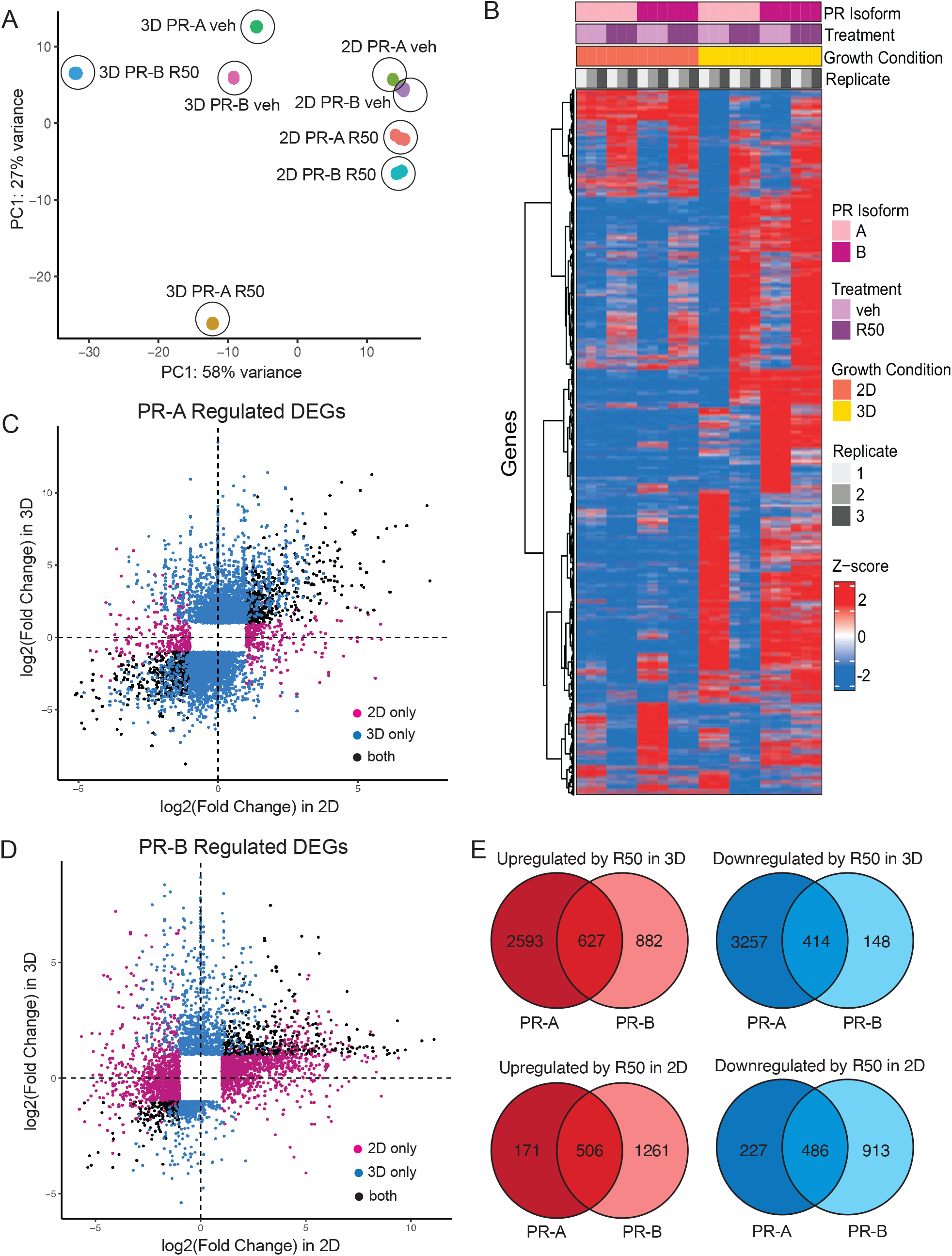
PR-A and PR-B control discrete transcriptomes. A. PCA Plot demonstrates that R5020 treatment and growth condition drive differential gene expression between RNA-seq samples. B. Clustered heatmap shows differential gene expression patterns in response to R5020 treatment when PR-A and PR-B expressing cells are grown in 2D and 3D conditions. All DEGs (log2Fold change > 1 or < -1; p.adjust < 0.05) C. Scatter plot shows that PR-A expressing cells have a larger response to R5020 treatment in 3D growth conditions. D. Scatter plot shows that PR-B expressing cells have a larger response to R5020 treatment in 2D growth conditions. E. Venn diagrams show the overlap between R5020-induced differentially expressed gene lists in PR-A and PR-B expressing cells.

We next quantified the magnitude of gene expression changes in response to R5020 treatment in both 2D and 3D conditions and plotted them as scatter plots for each PR isoform. In PR-A expressing cells the transcriptional response was more pronounced in 3D culture conditions, represented by the increase in differentially expressed genes (DEGs) denoted as blue dots (**Figure 2C**) with some DEGs being regulated in both 2D and 3D denoted as black dots. In contrast, PR-B expressing cells exhibited a greater response with more DEGs in 2D conditions marked as pink dots (**Figure 2D**). Consistent with this, Venn diagram analysis of DEGs revealed limited overlap between PR-A and PR-B transcriptional targets, underscoring the distinct regulatory programs driven by each PR isoform (**Figure 2E**). This large difference in gene regulation by PR-A and PR-B between *in vitro* growth conditions suggests that PR isoform-specific activity is likely influenced by the *in vivo* cellular microenvironment.

To explore the functional implications of PR isoform-specific gene regulation, we performed comparative gene set enrichment analysis (GSEA) using GO ontology terms on all differentially DEGs in PR-A and PR-B-expressing cells. This analysis revealed that PR-A and PR-B modulate distinct biological pathways and cellular functions depending on the growth conditions, highlighting their divergent transcriptional programs (**Figure 3A, Supplemental Table 1**). Specifically, PR-A-regulated DEGs were negatively enriched for pathways associated with cell cycle regulation and mitotic control. GSEA analysis of the PR-A R5020-induced DEGs in 3D using the MSigDB Hallmarks gene set further emphasized PR-A regulation of the cell cycle, with E2F target genes and G2M checkpoint genes among the top enriched hallmarks (**Figure 3B, Supplemental Table 2**). In contrast, PR-B-regulated genes were positively enriched for GO pathways related to cellular metabolic signaling and transmembrane receptor activity (**Figure 3A**). Further analysis of the PR-B R5020-induced DEGs in 2D using the MSigDB Hallmarks gene set showed that MYC targets were significantly upregulated, (**Figure 3C, Supplemental Table 3**), supporting a pro-growth and metabolic reprogramming role for PR-B.

**Figure 3.**
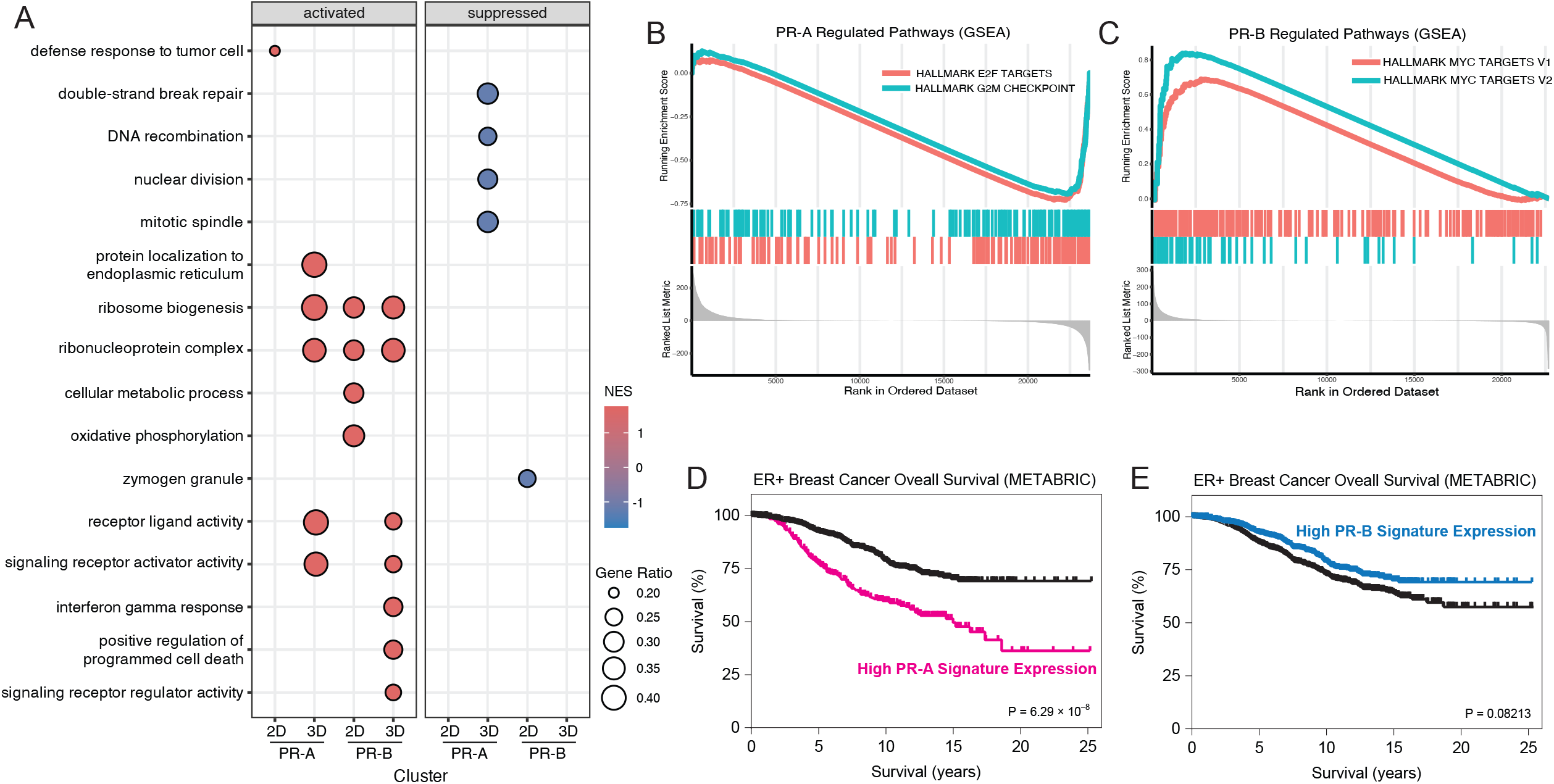
PR-A and PR-B sustain diverse transcriptomic landscapes. A. GSEA analysis dotplot shows that PR-A and PR-B regulate different pathways and cellular functions in 2D and 3D growth conditions. The top 8 enriched GO pathways are shown (FDR < 0.05). Dot color indicates normalized enrichment score (NES) for each gene set. Dot size indicates gene ratio for each gene set. B. GSEA enrichment plot demonstrates specific negative enrichment of cell cycle regulation, regulation of mitosis, and epithelial differentiation pathways within PR-A- regulated R50-induced DEGs in 3D growth conditions. C. GSEA enrichment plot demonstrates specific positive enrichment of cellular metabolic signaling, transmembrane signaling receptor activity, and MYC-driven proliferation within a PR-B-regulated R50-induced DEGs in 2D growth conditions. D. PR-A regulated gene signature is predictive of worse overall survival for patients in the METABRIC cohort. E. PR-B regulated gene signature is not predictive of better overall survival for patients in the METABRIC cohort.

To determine the clinical relevance of these PR isoform-specific gene signatures, we used the top 100 upregulated protein-coding DEGs (**Supplemental Table 4**) for each isoform and tested their prognostic value with a survival analysis using the METABRIC breast cancer cohort (31). A PR-A-regulated gene signature was significantly predictive of decreased overall survival (**Figure 3D**). Conversely, the PR-B-regulated gene signature did not correlate with patient outcomes (**Figure 3E**). These data suggest that a high PR A:B ratio may be associated with more aggressive disease and worse outcomes for breast cancer patients.

### PR-A and PR-B exhibit distinct genomic binding patterns

To investigate whether differences in transcriptional regulation arise from isoform-specific chromatin binding, we performed CUT&RUN to profile PR-A and PR-B genomic occupancy in response to R5020 treatment in 3D conditions. Peak overlap analysis revealed that the binding patterns of PR isoforms respond differently to R5020 treatment (**Figure 4A**). PR-A has 2,096 high-confidence peaks in the vehicle-treated condition, and shifts significantly to occupy 2,159 new sites after R5020 treatment with 1,805 remaining stable. In contrast, PR-B occupies only 81 unique binding sites in the absence of ligand and gains 1,356 new binding sites in the presence of R5020 while maintaining 947. Analysis of CUT&RUN signal intensities demonstrated that the majority of PR peaks are isoform-specific, with limited shared binding sites between PR-A and PR-B (less than 10%). Density plots demonstrated minimal overlap in CUT&RUN signal between PR-A and PR-B occupied regions (**Figure 4B**). Together these results suggest that PR-A is capable of more genomic binding in its unliganded state, while more PR-B genomic activity is dependent on ligand.

**Figure 4.**
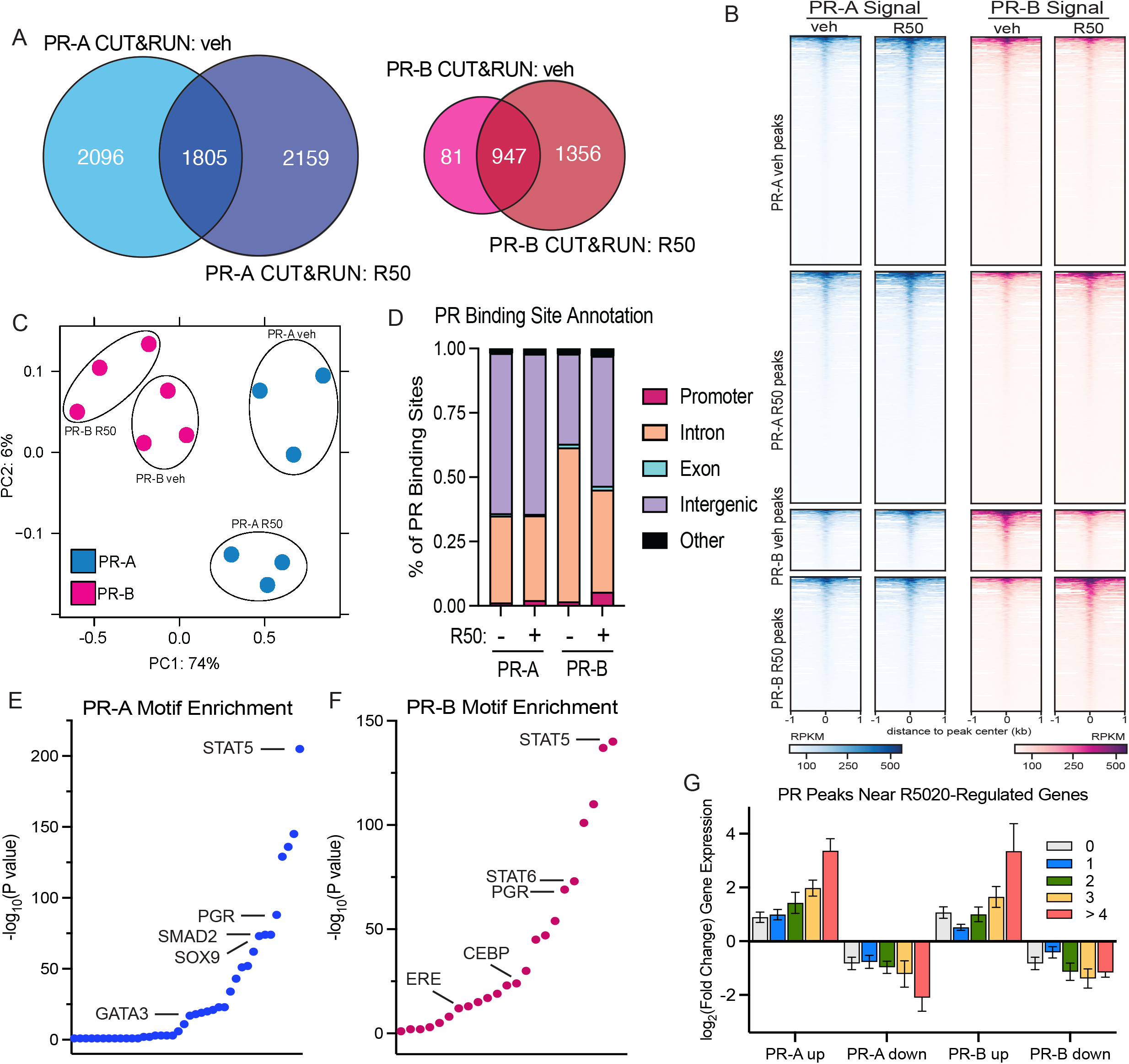
PR-A and PR-B have distinct genomic binding patterns. A. Venn diagrams represent peak overlap analysis for PR-A and PR-B in response to R5020 treatment. B. Density plots show limited overlap in CUT&RUN signal between PR-A occupied binding sites and PR-B occupied binding sites. C. PCA plot demonstrates that PR isoform is the main driver of differential binding patterns between CUT&RUN samples. Each data point represents one biological replicate (n = 3 per treatment group). D. Distribution of PR binding sites annotated to promoters, introns, exons, and intergenic regions. E. Homer motif analysis highlights the top transcription factor motifs enriched near PR-A binding sites. F. Homer motif analysis highlights the top transcription factor motifs enriched near PR-B binding sites. G. Average log2(FoldChange) of DEGs with 0,1, 2, 3 or >4 PR binding sites within 10kb of the TSS.

PCA of global PR binding sites showed that PR isoform identity was the primary driver of differential chromatin occupancy, further reinforcing the distinct binding profiles of PR-A and PR-B (**Figure 4C**). Annotation of PR binding sites to genomic features (**Figure 4D**) indicated differences in localization, with a notably greater proportion of PR-A binding occurring in intergenic regions, farther from the nearest transcriptional start site TSS. In contrast, more PR-B binding sites occur in promoter intronic and exonic regions. Motif analysis revealed that distinct transcription factor motifs were enriched at PR-A and PR-B binding sites, suggesting differential co-regulatory mechanisms that may contribute to the observed differences in transcriptional output (**Figure 4E**). Together, these data establish that PR-A and PR-B regulate divergent transcriptional programs by engaging distinct chromatin landscapes, emphasizing the isoform-specific mechanisms by which PRs influence biology. Analysis of PR binding site proximity to DEGs demonstrated that PR-A and PR-B target genes differ in their direct regulation by PR binding, with PR-B-regulated genes more likely to have proximal binding events (**Figure 4G**).

### PR-A and PR-B drive differential tumor growth in MIND xenograft models

To examine the *in vivo* relevance of PR isoform-specific functions, we utilized the mammary intraductal (MIND) tumor xenograft model (32,33) (**Figure 5A**). For these experiments we tagged T47D PR-A and PR-B cells with a luciferase reporter to visualize the growth of tumors over the course of their development. Bioluminescence imaging revealed that PR-A- and PR-B-expressing tumors exhibited distinct growth kinetics, with PR-B-expressing tumors displaying slightly more rapid expansion (**Figure 5B–C**); however, there was no significant difference in the final tumor volume. At 8 weeks post-tumor injection, animals were sacrificed, blood was collected and circulating tumor cells (CTCs) were expanded as soft agar colonies for quantification. We observed a significantly higher number of CTCs in the blood from animals carrying PR-A-expressing tumors, suggesting increased stem-like characteristics and metastatic potential (**Figure 5D**). Histological analysis of tumor sections showed that PR-B-expressing tumors were characterized by increased Ki67 staining, indicative of enhanced proliferation (**Figure 5E–F**). These *in vivo* results mirror the results of our *in vitro* phenotypic assays, where PR-A-expressing cells exhibited increased dissemination of breast tumor cells with stem-cell-like properties and PR-B-expressing cells showed increased proliferation within the primary tumor.

**Figure 5.**
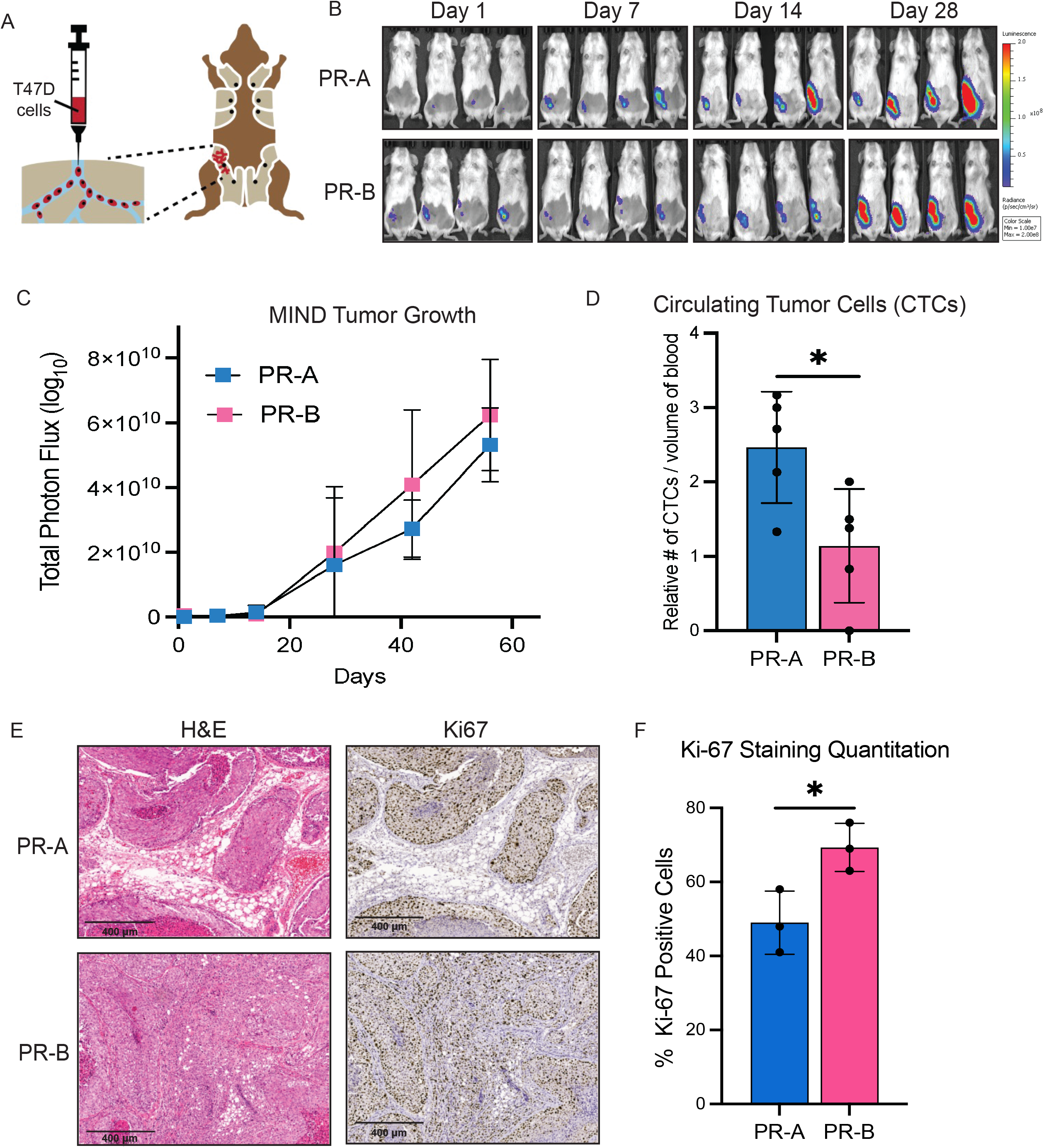
PR-A and PR-B drive differential growth patterns in MIND tumor xenograft models. A. Diagram illustrates mammary intraductal injection technique. B. Bioluminescent imaging of MIND xenograft tumors from injection of PR-A- and PR-B-expressing cells after 1, 7,14, and 28 days of growth (4 of 8 total mice per group shown). C. Quantification of PR-A and PR-B tumor growth over 60 days with bioluminescent imaging plotted as total proton flux over time. D. Quantification of circulating tumor cells (CTCs) per 100 µl blood recovered from mice injected with PR-A- or PR-B-expressing cells (n = 4). E. Representative images of H&E and Ki67 staining in PR-A and PR-B tumors. F. Quantification of total Ki67 staining in PR-A and PR- B expressing tumors (n =3).

## DISCUSSION

Progesterone receptor (PR) signaling is now increasingly recognized as a critical modulator of estrogen receptor-positive (ER+) luminal breast cancer biology. However, the distinct contributions of PR isoforms, PR-A and PR-B, to tumor progression and therapy resistance have remained largely unexplored. Our study provides novel insights into the isoform-specific functions of PR-A and PR-B by leveraging an isogenic model system that allows for exclusive expression of each isoform. Through transcriptomic, genomic, and functional analyses in both 2D and 3D growth conditions, we demonstrate that PR-A and PR-B regulate distinct gene expression programs, genomic landscapes, and tumorigenic phenotypes. These findings challenge the long-standing view of PR as a passive downstream effector of ER and further establish isoform-specific PR functions as key players in breast cancer heterogeneity.

Our results show that PR-A and PR-B exert divergent effects on breast cancer cell growth and plasticity, depending on the growth context (**Figure 1**). In 2D adherent conditions, PR-B-expressing cells displayed enhanced proliferation, consistent with PR-B’s role in driving MYC-associated metabolic signaling and mitogenic pathways. In contrast, 3D mammosphere assays revealed that PR-A promotes a more stem-like phenotype, supporting previous reports linking PR-A to breast cancer cell plasticity and endocrine resistance. This suggests that PR-A may contribute to tumor heterogeneity and therapy resistance (9-11) by maintaining a population of cancer cells with increased self-renewal capacity. Our PR-A gene signature obtained from breast cancer cells maintained in 3D conditions supports these findings.

These *in vitro* observations were recapitulated *in vivo* using the mammary intraductal (MIND) xenograft model (**Figure 5**). While PR-B-expressing tumors exhibited greater proliferative capacity, as evidenced by increased Ki67 staining of primary tumor cells, PR-A-expressing tumors were associated with greater breast cancer cell dissemination as measured by higher circulating tumor cell (CTC) counts, suggesting an enhanced propensity for survival in the circulation and ultimately metastatic potential. This aligns with our transcriptomic findings that PR-A-enriched gene signatures were negatively associated with epithelial differentiation and cell cycle regulation pathways, further supporting the hypothesis that PR-A promotes a less differentiated, more aggressive tumor state.

CUT&RUN profiling (**Figure 4**) provided additional mechanistic insight into how PR-A and PR-B differentially engage the chromatin landscape. PR-A exhibited more extensive genomic occupancy in the absence of ligand, suggesting a greater capacity for basal chromatin interactions, while PR-B binding was largely ligand-dependent. Interestingly, PR-B binding sites were more likely to occur proximal to transcription start sites (TSSs), whereas PR-A binding occurred more frequently in intergenic regions, potentially reflecting differences in their ability to engage distal enhancers. This observation suggests that PR isoforms recruit distinct transcriptional co-regulators, which could have implications for therapeutic targeting in an isoform selective manner.

Our transcriptomic analyses (**Figure 2, 3**) revealed striking differences in PR-A and PR-B-mediated gene regulation, with minimal overlap in DEGs between the isoforms. Gene set enrichment analysis (GSEA) (**Figure 3B-C**) further reinforced this functional divergence, revealing that PR-B activation drives positive enrichment of cellular metabolic processes and MYC-driven transcriptional programs, while PR-A activation resulted in negative enrichment of epithelial differentiation pathways, supporting a role in dedifferentiation and cellular plasticity. These differences may underlie the clinical observation that high PR-A:PR-B ratios correlate with poor outcomes in ER+ breast cancer (26), as PR-A-driven transcriptional programs may promote a more aggressive disease phenotype (Fig 3D). Our analysis of the METABRIC cohort demonstrated that the PR-A-driven gene signature correlates with worse overall survival, whereas the PR-B-driven signature had no significant prognostic value. This suggests that PR-A may contribute to endocrine therapy resistance, a hypothesis supported by our observation that PR-A is enriched in pathways linked to cell cycle deregulation and cellular plasticity.

Together, our findings provide a comprehensive functional and molecular characterization of PR isoform-specific activity in ER+ breast cancer cells. By demonstrating that PR-A and PR-B drive distinct phenotypic, transcriptional, and chromatin remodeling programs, our study challenges the conventional view of PR as a homogeneous entity and underscores the need for isoform-specific functional studies. A novel aspect of our work herein is the analysis of PR isoform-specific transcriptomes and cistromes performed in both traditional 2D as well as 3D mammosphere conditions. Modeling breast cancer in 3D culture systems more accurately recapitulates the architecture, cell–cell interactions, and microenvironmental cues present in tumors in vivo. Compared to traditional 2D monolayers, 3D cultures better reflect tissue polarity, drug responsiveness, and stem-like cell behavior, making them a more physiologically relevant model for studying hormone receptor signaling and therapy resistance. The use of 3D mammosphere cultures has revealed novel aspects of PR-A signaling that may provide useful biomarkers of early dissemination. Future studies should focus on further delineating the co-regulatory networks associated with PR isoforms of breast cancer cells in suspension, as well as investigating how PR phosphorylation and post-translational modifications influence their differential activities. Additionally, targeting PR isoform-specific mechanisms using novel PR modulators or chromatin-targeting therapies represents an exciting avenue for improving treatment efficacy in PR+ breast cancer.

## METHODS

### Cell Culture and Hormone Treatments

T47D cells were cultured in MEM (Corning) containing 5% FBS (Corning), 1% penicillin streptomycin (Gibco), nonessential amino acids (Gibco), and 6 ng/mL of insulin (Gibco). For experiments with R5020 (PerkinElmer) treatment, cells were hormone-starved in phenol red-free MEM (Gibco) containing 5% dextran-coated charcoal-stripped FBS (HyClone), 1% penicillin streptomycin (Gibco), nonessential amino acids (Gibco), and 6 ng/mL of insulin (Gibco) for 24 hours prior to treatment. For T47D PR isoform-specific models: T47D (Y; naturally occurring PR-null variant) (29) cells were transduced with lentiviral pLenti CMV Neo containing PR-A or PR-B (10). Cells were selected and maintained with 0.5 mg/ml G418 (Corning). All cell lines were cultured at 37°C under 5% CO_2_ in water-jacketed incubators. Cell lines were routinely tested for mycoplasma contamination with the E-myco Plus mycoplasma detection kit (Bulldog Bio, Inc). Cell lines were authenticated by ATCC, and results were compared with the ATCC short-tandem repeat (STR) database.

### Immunoblotting

Total protein was extracted from cells using radioimmunoprecipitation assay (RIPA) lite lysis buffer (150 mM NaCl, 6 mM Na2HPO4, 4 mM NaH2PO4, 2 mM EDTA, 100 mM NaF, 1% Triton-X 100, 1x complete mini protease inhibitors (Roche, 11836153001), and 1x PhosSTOP (Roche, 14 4906837001), supplemented with 1 mM PMSF, 5 mM NaF, 0.05 mM Na3VO4, 25 mM B15 glycerophosphate, and 20 mg/mL aprotinin. Protein concentration was quantified with Pierce BCA Protein Assay kit (ThermoScientific). 50 ug of total protein was fractionated by 8% SDS-PAGE and transferred to PVDF membranes (Millipore). Membranes were blocked for 1 hour at room temperature in phosphate-buffered saline/0.1% Tween-20 (PBS-T) containing 5% dried milk. Western blots were probed overnight at 4°C in PBS-T containing 1% milk using the following primary antibodies: PR-A/B (Santa Cruz Biotechnology Cat# sc-7208), alpha-tubulin (Cell Signaling, 3873), custom-made phospho-Ser294 PR antibody (Carol A. Lange - University of Minnesota Cat # CALange_PR294_8508) was commissioned from Thermo Fisher Scientific (26577046, 28412963). Secondary antibodies (HRP-conjugated goat anti-rabbit (Bio-Rad Cat # 170-6515) and goat anti-mouse (Bio-Rad Cat # 170-6516)) were used to detect their respective primary antibodies, and immunoreactive proteins were visualized on blue autoradiographic film (USA Scientific) following ECL detection with Super Signal West Pico Maximum Sensitivity Substrate (Pierce).

### Proliferation Assay

Live cell imaging was used to assess the growth kinetics of T47D CO, PR-A, and PR-B expressing cell lines. 10,000 cells were plated per well in 12-well plates. After 24 hours in phenol red–free, charcoal-stripped media, media were supplemented with either 10 nM R5020 or equivolume vehicle (ethanol). Brightfield images were taken every 24 hours with a BZ-X800 Fluorescent Microscope (Keyence), and cell numbers were quantified with BZ-X Series software (Keyence). Three independent experiments were performed with four technical replicates per condition per experiment.

### Mammosphere Assay

Mammosphere assays were performed in 24 well ultra-low attachment plates (Corning). Single cells were plated at a density of 1,000 cells per well and grown in a serum-free DMEM/F12 phenol-free medium (Corning) containing a final concentration of 1% methylcellulose (Sigma-Aldrich), 1x charcoal-stripped B27 proprietary supplement (Gibco), 1% penicillin-streptomycin, 5 mg/mL insulin (Gibco), 1 ng/mL hydrocortisone (Sigma-Aldrich, H0888), 20 ng/mL EGF, and 100 mM beta mercaptoethanol (MP Biomedicals). Mammospheres were allowed to grow for 8 days. To generate secondary mammospheres, primary spheres were collected and dissociated enzymatically with Accutase cell dissociation reagent (Gibco). Single cells were counted and replated in the DMEM/F12 methylcellulose containing media. The secondary mammospheres were allowed to grow for an additional 10 to 12 days. Mammospheres were imaged and quantified on a BZ-X800 Fluorescent Microscope (Keyence). Three independent experiments were performed with four technical replicates per condition per experiment.

### Soft Agar Colony Formation Assay

Cells were seeded in 6 well plates in sterile low-melting point agarose (Life Technologies) containing 5% charcoal-stripped FBS and the appropriate hormone treatment as indicated. Soft agar assays were allowed to proceed for 21 days at 37 °C. Afterward, cell colonies were fixed and stained in PBS containing 4% formaldehyde and 0.005% crystal violet. Colonies were quantitated using Fiji software. Three independent experiments were performed with three technical replicates per condition per experiment.

### RNA-seq

#### Sample collection and Sequencing

For 2D samples, T47D PR-A- or PR-B-expressing cells were hormone-starved in phenol red-free modified MEM media (Gibco) containing 10% dextran-coated charcoal-stripped FBS (HyClone) for 24 hours, then treated with 10nM R5020 or vehicle for 6 hours prior to sample collection. For 3D samples, T47D PR-A- or PR-B-expressing cells were plated in mammosphere conditions in 100 mm ultra-low attachment dishes with media containing charcoal-stripped B27 supplement and either 10nM R5020 or vehicle (ethanol). Mammospheres were allowed to grow for 6 days prior to sample collection. Three biological replicates per condition were collected. 1 µg of total RNA was extracted and purified using RNeasy Plus Kit (Qiagen) according to manufacturer’s protocol. This was repeated to collect a total of 3 biological replicates per condition. Purity of the total RNA samples was assessed via BioAnalyzer (Agilent) and samples with an RNA integrity score > 8 were used for library construction by the UMN Genomics Core. Libraries were built from individual samples with the TruSeq Stranded mRNA kit (Illumina). RNA-Seq libraries were pooled and sequenced on the Illumina NovaSeq X Plus with 50 bp paired-end reads.

#### Data Analysis

Quality scores across sequenced reads were assessed using FASTQC. All samples were high quality. Illumina adapters were removed using Trim-Galore. For alignment and transcript assembly, the sequencing reads were mapped to hg38 using STAR (34). Sorted reads were counted using HTSeq (35) and differential expression analysis was performed using DESeq2 (36). Genes with a p-value of <0.05 and a log_2_ fold change greater than 1 or less than - 1 were considered differentially expressed. Comparative gene set enrichment analysis (GSEA) was performed using the ClusterProfiler package (37) to identify differentially enriched pathways between experimental conditions.

### CUT&RUN

#### Sample collection and Sequencing

CUT&RUN (38) was performed using T47D PR-A- or PR-B-expressing cells cultured in mammosphere conditions in 100 mm ultra-low attachment dishes with media containing charcoal-stripped B27 supplement (Gibco) and either 10nM R5020 or vehicle (ethanol). Mammospheres were allowed to grow for 6 days prior to sample collection. Mammospheres were then harvested, washed, separated into a single-cell suspension using Accutase (STEMCELL Technologies), and bound to activated Concanavalin A coated magnetic beads (Epicypher). Three biological replicates per condition were collected. Cells were permeabilized with Wash buffer (20mM HEPES pH 7.5, 150mM NaCl, 0.5 mM spermidine, 0.01% digitonin) and incubated with the indicated antibody at 4°C with constant agitation overnight. Cells were washed twice more before incubation with recombinant p-AG MNase (Epicypher) at 4°C for 2 hours. Liberated DNA was purified, and libraries were prepared using the NEB Ultra II DNA Library Kit (New England Biolabs) and amplified with 14 cycles of PCR. Amplified libraries were purified with AMPure beads (Beckman Coulter), quantified via Qubit (Life Technologies), and quality was assessed using the BioAnalyzer (Agilent) High-Sensitivity DNA kit. CUT&RUN libraries were pooled and sequenced on the Illumina NextSeq 2000 with 50 bp paired-end reads in the UMN Genomics Core.

#### Data Analysis

Quality scores across sequenced reads were assessed using FASTQC. Illumina adapters were removed using Trim Galore (39). Paired-end reads were mapped to hg38 using Bowtie2 (40), and peaks were called using MACS2 (41). Consensus peak sets for downstream analysis were derived using IDR (42) using three replicates per target and a cut-off of 0.05. DiffBind (43) was used to assess sample quality and differential PR binding. Deeptools (44) was used for visualizing CUT&RUN signal over genomic regions. HOMER (45) was used for motif analysis and peak annotation.

### Mammary Intraductal (MIND) Induction of PR-expressing Tumors

Six-week-old female NOD Cg-*Prk-dc*^*scid*^ *Il2rg*^*tm1Wj l*^/SzJ (NSG) mice (Jackson Laboratories) were housed 4 per cage in UMN Research Animal Resources (RAR) facilities and allowed to acclimate for 1 week prior to handling. Prior to injections, T47D PR-A and T47D PR-B cell lines (10) were transduced with lentiviral vector pLenti CMV Puro LUC (w168-1) (Addgene plasmid #17477) and luciferase activity was confirmed with a luciferase activity assay kit (Promega). The luciferase vector was maintained with 0.5 μg/mL puromycin selection.

For each injection into a single mouse, 24 hour in vitro homotypic cell clusters were made with the T47D PR-A-luc or T47D PR-B-luc cell lines in ultra-low attachment plates (Corning) using 15,000 cells/well/mouse and Mammocult media (Stem Cell Technologies) according to manufacturer’s instructions. Cell clusters were injected into the mammary ducts of mammary fat pad 9 (MFP9) as described (46). Briefly, animals that had been previously dehaired around the nipple area were anesthetized with 2-5% inhaled isoflurane, the area was cleansed with ethanol, the nipple of MFP9 was cut off with 4 mm Vannas spring scissors, and 10 μL of PBS plus cell clusters was injected into the ductal opening with a 30-gauge Hamilton syringe with a ½ inch blunt-ended needle. The area was swabbed with iodine and animals were allowed to recover under a heating pad. Animals were given ad-libitum access to water containing 2 μM 17 β-estradiol after injection and for the duration of the experiment.

Monitoring of luciferase expression was performed on an IVIS Spectrum (PerkinElmer) using Living Image software (Caliper LifeSciences) in the UMN Imaging Centers. Images were taken 1 day, 1 week, 2 weeks, 4 weeks, 6 weeks, and 8 weeks post-intraductal injection using 100 μL of Rediject D-luciferin (Revvity) injected intraperitoneally. Quantitation of photon emittance was also performed with Living Image software. Animal protocols for this study were approved by the UMN Institutional Animal Care and Use Committee (IACUC). Two independent experiments were performed, with a total of 8 animals per group.

### Detection of Circulating Tumor Cells (CTCs)

Circulating tumor cells (CTCs) were isolated at the end of the MIND tumor growth experiment. Four tumor-bearing mice per group were anesthetized, and blood was collected by cardiac puncture. Blood elements and tumor cells were isolated by density centrifugation using Isolymph (CTL Scientific Supply Corp). Blood (500-800 µL) was diluted with 3 mL DMEM supplemented with 5% FBS, overlaid gently on a 2 mL cushion of Isolymph and centrifuged at 1700 RPM for 30 min at 4°C with the brakes off. The buffy coat with cells was collected and washed once with DMEM supplemented with 5% FBS, and centrifuged at 1200 RPM for 10 min. The cell pellet was resuspended in 400 µL of media, mixed with 600 µL of 0.8% agarose, and plated on a bottom layer of 0.8% agar solidified in the bottom of a 6 well plate. Colonies were quantified after 21 days.

### Immunohistochemistry (IHC)

Three tumors per group were removed and fixed in 4% fresh paraformaldehyde for 24 hrs and then maintained in 70% ethanol until they were ready for paraffin embedding and sectioning by the UMN CTSI Histology lab. Unstained tumor sections (4 µm) were de-paraffinized and rehydrated using standard methods. For antigen retrieval, slides were incubated in 6.0 pH buffer (Reveal Decloaking reagent, Biocare Medical, Concord, CA) in a steamer for 30 min at 95-98°C, followed by a 20 min cool down period. Endogenous peroxidase activity was quenched by slide immersion in 3% hydrogen peroxide solution (Peroxidazed, Biocare Medical, Concord, CA) for 10 min followed by TBS-Tween (TBST) rinse. A serum-free blocking solution (Sniper, Biocare Medical, Concord, CA) was placed on sections for 15 min. Blocking solution was removed and slides were incubated in primary antibody diluted in 10% blocking solution/90% TBST. Mouse monoclonal anti-Ki67 (Dako, Carpentaria, CA 1:200) was applied and incubated for 60 min at room temperature followed by TBST rinse and detection with Novalink polymer (Leica Biosystems, Deer Park, IL) according to manufacturer’s instructions at room temperature. All slides then proceeded with TBST rinse and visualization with diaminobenzidine (DAB) (Biolegend, Dedham, MA). Slides were incubated for 5 min followed by TBS rinse then counterstained with CAT Hematoxylin (Biocare, Concord, CA) for 5 min. Slides were then dehydrated and coverslipped.Total Ki67 across the entire slide was quantified using QuPath.

### Statistical Analysis

The data were tested for a normal distribution using the Shapiro-Wilks normality test and the homogeneity of variances using the Bartlett test. After confirming that the data met these two requirements, statistical analyses were performed. For comparisons between more than two groups, either one-way or two-way ANOVA along with the Tukey multiple comparison test was used for mean values. For comparisons between two groups, the Student t test was used. Significance was determined at a 95% confidence level.

## DATA ACCESS

All raw and processed next generation sequencing data associated with this manuscript is available for download from the NCBI Gene Expression Omnibus repository under accession codes GSE293884 and GSE293885. Bash and R scripts for analysis of RNA-seq and CUT&RUN data is available in a dedicated github repository https://github.com/negillis/PR-Isoforms-Analysis.

## ACKNOWLEDGEMENTS

This work was supported by resources and staff at the University of Minnesota Genomics Core (UMGC), the University Imaging Centers (UIC), and the Clinical and Translational Science Institute’s (CSTI) Histology and Digital Imaging team.

We would like to acknowledge Drs. Carol Sartorius and Jessica Finlay-Schultz for their guidance and expertise on the analysis of this data.

## REFERENCES

1. Arpino, G., De Angelis, C., Giuliano, M., Giordano, A., Falato, C., De Laurentiis, M. and De Placido, S. (2009) Molecular mechanism and clinical implications of endocrine therapy resistance in breast cancer. Oncology, 77 Suppl 1, 23–37.

2. Li, Z., Wu, Y., Yates, M.E., Tasdemir, N., Bahreini, A., Chen, J., Levine, K.M., Priedigkeit, N.M., Nasrazadani, A., Ali, S. et al. (2022) Hotspot ESR1 Mutations Are Multimodal and Contextual Modulators of Breast Cancer Metastasis. Cancer Res, 82, 1321–1339.

3. Liang, J., Yao, X., Aouad, P., Wang, B.E., Crocker, L., Chaudhuri, S., Liang, Y., Darmanis, S., Giltnane, J., Moore, H.M. et al. (2025) ERalpha dysfunction caused by ESR1 mutations and therapeutic pressure promotes lineage plasticity in ER(+) breast cancer. Nat Cancer, 6, 357–371.

4. Bardou, V.J., Arpino, G., Elledge, R.M., Osborne, C.K. and Clark, G.M. (2003) Progesterone receptor status significantly improves outcome prediction over estrogen receptor status alone for adjuvant endocrine therapy in two large breast cancer databases. J Clin Oncol, 21, 1973–1979.

5. Mohammed, H., Russell, I.A., Stark, R., Rueda, O.M., Hickey, T.E., Tarulli, G.A., Serandour, A.A., Birrell, S.N., Bruna, A., Saadi, A. et al. (2015) Progesterone receptor modulates ERalpha action in breast cancer. Nature, 523, 313–317.

6. Singhal, H., Greene, M.E., Tarulli, G., Zarnke, A.L., Bourgo, R.J., Laine, M., Chang, Y.F., Ma, S., Dembo, A.G., Raj, G.V. et al. (2016) Genomic agonism and phenotypic antagonism between estrogen and progesterone receptors in breast cancer. Sci Adv, 2, e1501924.

7. Daniel, A.R., Gaviglio, A.L., Knutson, T.P., Ostrander, J.H., D’Assoro, A.B., Ravindranathan, P., Peng, Y., Raj, G.V., Yee, D. and Lange, C.A. (2015) Progesterone receptor-B enhances estrogen responsiveness of breast cancer cells via scaffolding PELP1- and estrogen receptor-containing transcription complexes. Oncogene, 34, 506–515.

8. Singhal, H., Greene, M.E., Zarnke, A.L., Laine, M., Al Abosy, R., Chang, Y.F., Dembo, A.G., Schoenfelt, K., Vadhi, R., Qiu, X. et al. (2018) Progesterone receptor isoforms, agonists and antagonists differentially reprogram estrogen signaling. Oncotarget, 9, 4282–4300.

9. Truong, T.H., Dwyer, A.R., Diep, C.H., Hu, H., Hagen, K.M. and Lange, C.A. (2019) Phosphorylated Progesterone Receptor Isoforms Mediate Opposing Stem Cell and Proliferative Breast Cancer Cell Fates. Endocrinology, 160, 430–446.

10. Diep, C.H., Spartz, A., Truong, T.H., Dwyer, A.R., El-Ashry, D. and Lange, C.A. (2024) Progesterone Receptor Signaling Promotes Cancer Associated Fibroblast Mediated Tumorigenicity in ER+ Breast Cancer. Endocrinology, 165.

11. Dwyer, A.R., Truong, T.H., Kerkvliet, C.P., Paul, K.V., Kabos, P., Sartorius, C.A. and Lange, C.A. (2021) Insulin receptor substrate-1 (IRS-1) mediates progesterone receptor-driven stemness and endocrine resistance in oestrogen receptor+ breast cancer. Br J Cancer, 124, 217–227.

12. Anderson, E. (2002) The role of oestrogen and progesterone receptors in human mammary development and tumorigenesis. Breast Cancer Res, 4, 197–201.

13. Beral, V. and Million Women Study, C. (2003) Breast cancer and hormone-replacement therapy in the Million Women Study. Lancet, 362, 419–427.

14. Clarke, C.A., Glaser, S.L., Uratsu, C.S., Selby, J.V., Kushi, L.H. and Herrinton, L.J. (2006) Recent declines in hormone therapy utilization and breast cancer incidence: clinical and population-based evidence. J Clin Oncol, 24, e49–50.

15. Hofseth, L.J., Raafat, A.M., Osuch, J.R., Pathak, D.R., Slomski, C.A. and Haslam, S.Z. (1999) Hormone replacement therapy with estrogen or estrogen plus medroxyprogesterone acetate is associated with increased epithelial proliferation in the normal postmenopausal breast. J Clin Endocrinol Metab, 84, 4559–4565.

16. Ross, R.K., Paganini-Hill, A., Wan, P.C. and Pike, M.C. (2000) Effect of hormone replacement therapy on breast cancer risk: estrogen versus estrogen plus progestin. J Natl Cancer Inst, 92, 328–332.

17. Rossouw, J.E., Anderson, G.L., Prentice, R.L., LaCroix, A.Z., Kooperberg, C., Stefanick, M.L., Jackson, R.D., Beresford, S.A., Howard, B.V., Johnson, K.C. et al. (2002) Risks and benefits of estrogen plus progestin in healthy postmenopausal women: principal results From the Women’s Health Initiative randomized controlled trial. JAMA, 288, 321–333.

18. De Vivo, I., Huggins, G.S., Hankinson, S.E., Lescault, P.J., Boezen, M., Colditz, G.A. and Hunter, D.J. (2002) A functional polymorphism in the promoter of the progesterone receptor gene associated with endometrial cancer risk. Proc Natl Acad Sci U S A, 99, 12263–12268.

19. Johnatty, S.E., Spurdle, A.B., Beesley, J., Chen, X., Hopper, J.L., Duffy, D.L., Chenevix-Trench, G. and Kathleen Cuningham Consortium for Research in Familial Breast, C. (2008) Progesterone receptor polymorphisms and risk of breast cancer: results from two Australian breast cancer studies. Breast Cancer Res Treat, 109, 91–99.

20. Kotsopoulos, J., Tworoger, S.S., De Vivo, I., Hankinson, S.E., Hunter, D.J., Willett, W.C. and Chen, W.Y. (2009) +331G/A variant in the progesterone receptor gene, postmenopausal hormone use and risk of breast cancer. Int J Cancer, 125, 1685–1691.

21. Romano, A., Lindsey, P.J., Fischer, D.C., Delvoux, B., Paulussen, A.D., Janssen, R.G. and Kieback, D.G. (2006) Two functionally relevant polymorphisms in the human progesterone receptor gene (+331 G/A and progins) and the predisposition for breast and/or ovarian cancer. Gynecol Oncol, 101, 287–295.

22. Terry, K.L., De Vivo, I., Titus-Ernstoff, L., Sluss, P.M. and Cramer, D.W. (2005) Genetic variation in the progesterone receptor gene and ovarian cancer risk. Am J Epidemiol, 161, 442–451.

23. Graham, J.D. and Clarke, C.L. (2002) Expression and transcriptional activity of progesterone receptor A and progesterone receptor B in mammalian cells. Breast Cancer Res, 4, 187–190.

24. Graham, J.D., Yeates, C., Balleine, R.L., Harvey, S.S., Milliken, J.S., Bilous, A.M. and Clarke, C.L. (1995) Characterization of progesterone receptor A and B expression in human breast cancer. Cancer Res, 55, 5063–5068.

25. Graham, J.D., Yager, M.L., Hill, H.D., Byth, K., O’Neill, G.M. and Clarke, C.L. (2005) Altered progesterone receptor isoform expression remodels progestin responsiveness of breast cancer cells. Mol Endocrinol, 19, 2713–2735.

26. Elia, A., Saldain, L., Vanzulli, S.I., Helguero, L.A., Lamb, C.A., Fabris, V., Pataccini, G., Martinez-Vazquez, P., Burruchaga, J., Caillet-Bois, I. et al. (2023) Beneficial Effects of Mifepristone Treatment in Patients with Breast Cancer Selected by the Progesterone Receptor Isoform Ratio: Results from the MIPRA Trial. Clin Cancer Res, 29, 866–877.

27. Bonneterre, J., Bosq, J., Jamme, P., Valent, A., Gilles, E.M., Zukiwski, A.A., Fuqua, S.A., Lange, C.A. and O’Shaughnessy, J. (2016) Tumour and cellular distribution of activated forms of PR in breast cancers: a novel immunohistochemical analysis of a large clinical cohort. ESMO Open, 1, e000072.

28. Knutson, T.P., Truong, T.H., Ma, S., Brady, N.J., Sullivan, M.E., Raj, G., Schwertfeger, K.L. and Lange, C.A. (2017) Posttranslationally modified progesterone receptors direct ligand-specific expression of breast cancer stem cell-associated gene programs. J Hematol Oncol, 10, 89.

29. Sartorius, C.A., Groshong, S.D., Miller, L.A., Powell, R.L., Tung, L., Takimoto, G.S. and Horwitz, K.B. (1994) New T47D breast cancer cell lines for the independent study of progesterone B- and A-receptors: only antiprogestin-occupied B-receptors are switched to transcriptional agonists by cAMP. Cancer Res, 54, 3868–3877.

30. Lange, C.A., Shen, T. and Horwitz, K.B. (2000) Phosphorylation of human progesterone receptors at serine-294 by mitogen-activated protein kinase signals their degradation by the 26S proteasome. Proc Natl Acad Sci U S A, 97, 1032–1037.

31. Curtis, C., Shah, S.P., Chin, S.F., Turashvili, G., Rueda, O.M., Dunning, M.J., Speed, D., Lynch, A.G., Samarajiwa, S., Yuan, Y. et al. (2012) The genomic and transcriptomic architecture of 2,000 breast tumours reveals novel subgroups. Nature, 486, 346–352.

32. Quinn, H.M., Battista, L., Scabia, V. and Brisken, C. (2024) Preclinical Mouse Intraductal Model (MIND) to Study Metastatic Dormancy in Estrogen Receptor-Positive Breast Cancer. Methods Mol Biol, 2811, 101–112.

33. Truong, T.H., Benner, E.A., Hagen, K.M., Temiz, N.A., Kerkvliet, C.P., Wang, Y., Cortes-Sanchez, E., Yang, C.H., Trousdell, M.C., Pengo, T. et al. (2021) PELP1/SRC-3-dependent regulation of metabolic PFKFB kinases drives therapy resistant ER(+) breast cancer. Oncogene, 40, 4384–4397.

34. Dobin, A., Davis, C.A., Schlesinger, F., Drenkow, J., Zaleski, C., Jha, S., Batut, P., Chaisson, M. and Gingeras, T.R. (2013) STAR: ultrafast universal RNA-seq aligner. Bioinformatics, 29, 15–21.

35. Putri, G.H., Anders, S., Pyl, P.T., Pimanda, J.E. and Zanini, F. (2022) Analysing high-throughput sequencing data in Python with HTSeq 2.0. Bioinformatics, 38, 2943–2945.

36. Love, M.I., Huber, W. and Anders, S. (2014) Moderated estimation of fold change and dispersion for RNA-seq data with DESeq2. Genome Biol, 15, 550.

37. Yu, G., Wang, L.G., Han, Y. and He, Q.Y. (2012) clusterProfiler: an R package for comparing biological themes among gene clusters. OMICS, 16, 284–287.

38. Meers, M.P., Bryson, T.D., Henikoff, J.G. and Henikoff, S. (2019) Improved CUT&RUN chromatin profiling tools. eLife, 8, e46314.

39. Krueger, F. 0.6.10 ed. The Babraham Institute.

40. Langmead, B. and Salzberg, S.L. (2012) Fast gapped-read alignment with Bowtie 2. Nat Methods, 9, 357–359.

41. Zhang, Y., Liu, T., Meyer, C.A., Eeckhoute, J., Johnson, D.S., Bernstein, B.E., Nusbaum, C., Myers, R.M., Brown, M., Li, W. et al. (2008) Model-based analysis of ChIP-Seq (MACS). Genome Biol, 9, R137.

42. Li, Q., Brown, J.B., Huang, H. and Bickel, P.J. (2011) Measuring reproducibility of high-throughput experiments. The Annals of Applied Statistics, 5, 1752–1779, 1728.

43. Ross-Innes, C.S., Stark, R., Teschendorff, A.E., Holmes, K.A., Ali, H.R., Dunning, M.J., Brown, G.D., Gojis, O., Ellis, I.O., Green, A.R. et al. (2012) Differential oestrogen receptor binding is associated with clinical outcome in breast cancer. Nature, 481, 389–393.

44. Ramirez, F., Ryan, D.P., Gruning, B., Bhardwaj, V., Kilpert, F., Richter, A.S., Heyne, S., Dundar, F. and Manke, T. (2016) deepTools2: a next generation web server for deep-sequencing data analysis. Nucleic Acids Res, 44, W160–165.

45. Heinz, S., Benner, C., Spann, N., Bertolino, E., Lin, Y.C., Laslo, P., Cheng, J.X., Murre, C., Singh, H. and Glass, C.K. (2010) Simple combinations of lineage-determining transcription factors prime cis-regulatory elements required for macrophage and B cell identities. Mol Cell, 38, 576–589.

46. Behbod, F., Kittrell, F.S., LaMarca, H., Edwards, D., Kerbawy, S., Heestand, J.C., Young, E., Mukhopadhyay, P., Yeh, H.W., Allred, D.C. et al. (2009) An intraductal human-in-mouse transplantation model mimics the subtypes of ductal carcinoma in situ. Breast Cancer Res, 11, R66.

